# Intrinsically disordered N-terminal regions suppress cotranslational protein degradation

**DOI:** 10.64898/2026.06.13.732044

**Authors:** Donghong Ju, Daniel Xie, Junzhe Wang, ShiChao Wu, Li Li, Youming Xie

## Abstract

Intrinsically disordered regions (IDRs) of proteins are thought to be inherently sensitive to proteolysis and considered one of the key components constituting an efficient degron. Here we report that IDRs can also suppress protein degradation. Our recent study showed that yeast ribosomal proteins, while posttranslationally stable, are subject to cotranslational protein degradation (CTPD). In mapping the degron responsible for CTPD of ribosomal protein Rpl8A, we found that its N-terminal IDR suppresses CTPD, whereas the adjacent structured domain acts as a degron. We further assessed the N-terminal IDRs of 9 other yeast proteins and found that they all inhibit CTPD. These results suggest that suppression of CTPD is likely a generic function of N-terminal IDRs. Moreover, we showed that the N-terminal IDR of human ribosomal protein hRpl7A also functions as a stabilizer against CTPD in human cells. When transplanted to the N-terminus of cystic fibrosis transmembrane conductance regulator (CFTR), the N-terminal IDR of hRpl7A reduces CTPD of CFTR by more than 80%. Thus, the stabilizer function of N-terminal IDRs is conserved from yeast to human. Using mass spectrometry, we demonstrated that HSP70 chaperone proteins Ssa and Ssb bind to the N-terminal IDR of Rpl8A. These data suggest that N-terminal IDRs may inhibit CTPD through recruiting HSP70 chaperone proteins to nascent chains, thereby facilitating cotranslational folding. Our study unveils a new role for IDRs in suppressing CTPD.

## Introduction

Intrinsically disordered regions (IDRs) are defined as polypeptide segments that are unable to acquire a stable tertiary (3D) structure (1–3). IDRs are highly abundant across the proteomes of all kingdoms of life (4–7). While widely present, the biological significance of IDRs was once a perplexing problem, caused by the early assumption that proteins exert their functions exclusively through molecular interactions mediated by structured domains. Recent studies showed that IDRs conduct a variety of biological functions such as transcription, cell signaling, formation of biomolecular condensates, and adaptation to extreme environments (8–12). Because of lacking a 3D structure, IDRs are thought to be inherently sensitive to proteolysis. In the process of proteasomal degradation, a substrate’s IDR engages with the ATPase ring of 19S regulatory proteasome (13, 14). This interaction is supposed to trigger substrate unfolding, an essential step for the substrate to enter the 20S core proteasome for degradation (15). Thus, IDRs are often considered a crucial element for an efficient degron.

A large fraction of nascent chains is subject to proteasomal degradation during synthesis, an event known as cotranslational protein degradation (CTPD) (16–20). The term CTPD is also used to describe the degradation of newly synthesized proteins already released from the ribosome but not yet folded. It has been suggested that CTPD serves as a quality control to eliminate defective nascent chains deemed unable to fold properly. This postulation was largely based on the studies using nascent chains incorporated with amino acid analogs or engineered polypeptides produced by translation of mRNAs lacking a stop codon or containing obstacles to translation elongation (18, 21, 22). To comprehend the biological significance of CTPD, we applied a systematic proteomic approach to identify native CTPD substrates (23). Intriguingly, approximately 60% of the yeast proteome encounters CTPD under normal growth conditions. Many proteins are subject to high level CTPD, with more than 30% of their nascent chains degraded cotranslationally. These results indicate that CTPD plays a role in gauging the steady-state levels of most endogenous proteins. Notably, we found that there is no correlation between CTPD and posttranslational protein stability (or protein half-life). One particularly interesting example is ribosomal proteins. While ribosomal proteins have long half-lives (24), all 62 ribosomal proteins collected by our proteomic analysis are subject to CTPD (25). These observations suggest that the degrons driving CTPD are masked in mature ribosomal proteins.

In this study, we set out to dissect the degron that mediates CTPD of yeast ribosomal protein Rpl8A. The CTPD degron is located within the first 100 amino acids that include an N-terminal IDR and a structured domain. Surprisingly, the structured domain acts as a degron, whereas the N-terminal IDR suppresses CTPD. Moreover, we showed that the N-terminal IDRs of 9 other yeast proteins all inhibit CTPD. The N-terminal IDR of human ribosomal protein hRpl7A also suppresses CTPD in human cells. We further applied proteomic analysis and AlphaFold-based modeling to elucidate the mechanism by which N-terminal IDRs suppress CTPD. Our study uncovers a new role for IDRs in suppressing CTPD. The implications of this finding are discussed.

## Results

### Mapping the degron of Rpl8A

We recently showed that the degron directing CTPD of Rpl8A resides within the N-terminal 100 amino acids (Rpl8A_1-100_) (25). Since Rpl8A is posttranslationally stable, the degron must be shielded in the mature protein. We found that a mutant deleted of C-terminal residues 201-256 is unstable posttranslationally, suggesting that the degron is masked by the C-terminal region via intramolecular interaction. Deletion of the C-terminal region likely leaves the degron exposed continuously, and therefore, it not only directs CTPD but also mediates posttranslational degradation. To further map the degron, we used AlphaFold to analyze the structure of Rpl8A obtained from the Protein Data Bank (26, 27). Rpl8A_1-100_ is composed of two distinct regions (Fig. 1A). The N-terminal region of the first 50 amino acids (Rpl8A_1-50_) is intrinsically disordered, whereas the adjacent region of residues 51-100 (Rpl8A_51-100_) contains 2 helices in contact with C-terminal domains. Secondary structure predictions, conducted by the JPrep server (28), also revealed that Rpl8A_1-50_ is unstructured, whereas Rpl8A_51-100_ possesses 2 helices (Fig. S1A). In addition, we plotted per-residue disorder for Rpl8A_1-100_ using the deep learning-based predictor Metapredict V2 (29). As shown in Fig. S1B, the residues in the N-terminal half of Rpl8A_1-100_ have a high probability of being disordered. Based on the structural analysis, both Rpl8A_1-50_ and Rpl8A_51-100_ could potentially serve as a degron. As mentioned, the degron may be masked in the mature protein via interactions with the C-terminal region involving residues 201-256. Rpl8A_51-100_ is at least in contact with residues 234, 236 and 237 (Fig. 1A). On the other hand, Rpl8A_1-50_ is an IDR, which is supposed to be sensitive to proteolysis.

**Figure 1.**
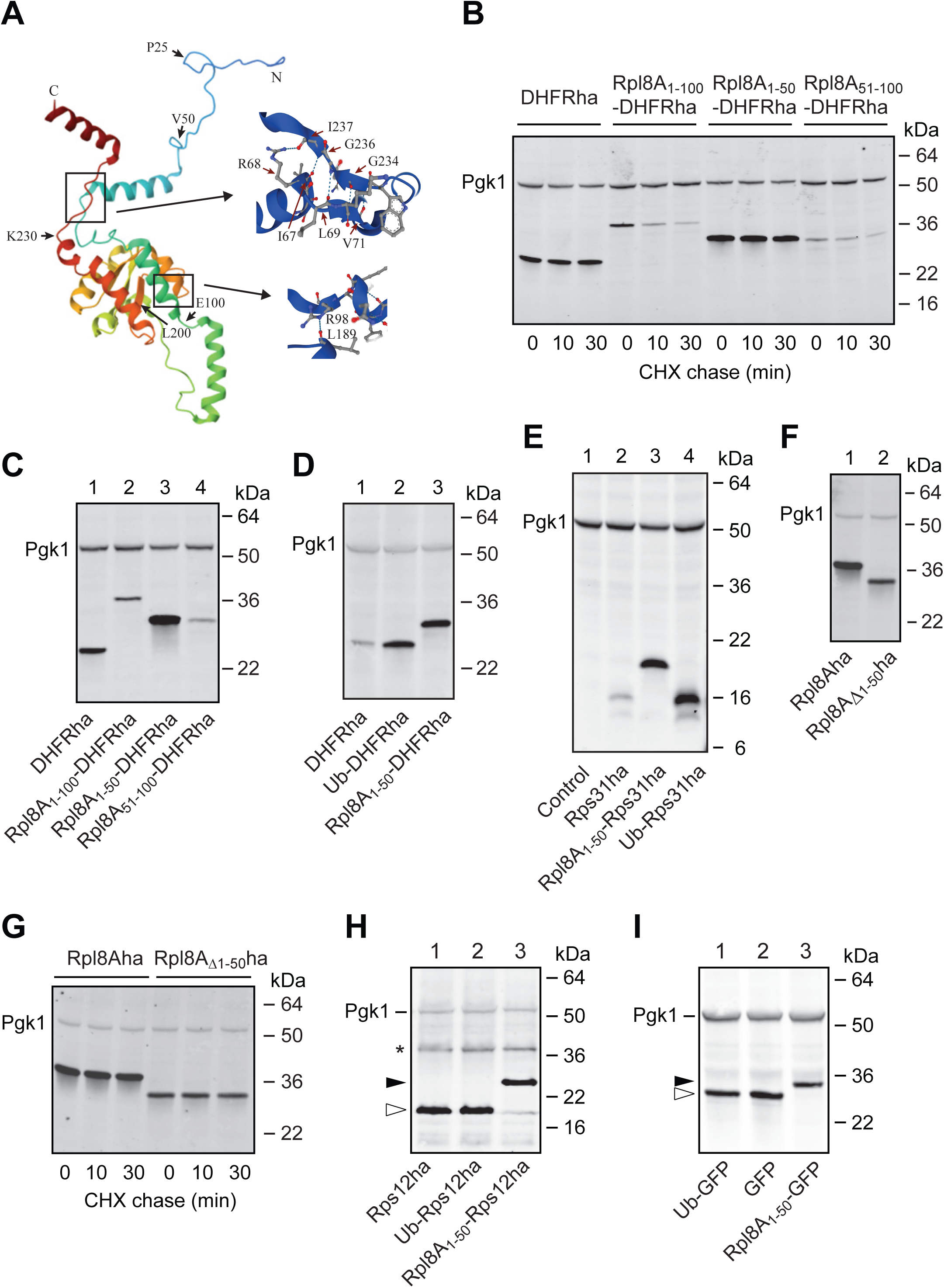
The N-terminal IDR of Rpl8A acts as a stabilizer against CTPD. (A) The N-terminal region of Rpl8A is an IDR. Shown here is a ribbon representation of Rpl8A structure obtained from the Protein Data Bank (PDB 4U51). Segments from N- to C-terminus are labeled with different colors (from indigo to red). The annotations of amino acid contacts involved in intramolecular interactions (enlarged) were obtained from AlphaFold analysis. (B) Measurement of degradation of DHFRha, Rpl8A_1-100_-DHFRha, Rpl8A_1-50_-DHFRha, and Rpl8A_51-100_-DHFRha by CHX-chase assay. Yeast cells expressing various proteins from a low-copy number vector under the control of *CUP1* promoter were harvested at 0, 10 and 30 min during CHX chase. Cell lysates were applied to Western blotting analysis with anti-ha and anti-Pgk1 antibodies. Pgk1 served as a loading control. (C) Comparison of the steady-state levels of DHFRha, Rpl8A_1-100_-DHFRha, Rpl8A_1-50_-DHFRha, and Rpl8A_51-100_-DHFRha. Anti-ha/anti-Pgk1 Western blotting was performed to analyze the crude extracts of yeast transformants as indicated. (D) Comparable inhibitory effects of N-terminal Rpl8A_1-50_ and Ub on CTPD of DHFRha. Yeast cells expressing DHFRha, Ub-DHFRha, and Rpl8A_1-50_-DHFRha were analyzed by Western blotting as described above. (E) Rpl8A_1-50_ protects Rps31 from CTPD. Western blotting analysis was conducted for yeast cells expressing Rps31ha, Rpl8A_1-50_-Rps31ha, and Ub-Rps31. A void vector served as a control. (F) Comparison of the steady-state levels of Rpl8Aha and Rpl8A_Δ1-50_ha by Western blotting with anti-ha and anti-Pgk1 antibodies. (G) Rpl8Aha and Rpl8A_Δ1-50_ha are posttranslationally stable. CHX chase analysis was performed to monitor posttranslational degradation of Rpl8Aha and Rpl8A_Δ1-50_-ha. (H) Western blotting analysis of yeast transformants expressing Rps12ha, Ub-Rps12ha, and Rpl8A_1-50_-Rps12ha. Open and filled arrowheads marked Rps12ha and Rpl8A_1-50_-Rps12ha, respectively. (I) Yeast transformants expressing GFP, Ub-GFP, and Rpl8A_1-50_-GFP were analyzed by Western blotting with anti-GFP and anti-Pgk1 antibodies. Open and filled arrowheads marked GFP and Rpl8A_1-50_-GFP, respectively.

We went on to examine if Rpl8A_1-50_ or Rpl8A_51-100_ or both could function as a degron. Our early work showed that the degron of Rpl8A is transplantable (25). When added to the N-terminus, Rpl8A_1-100_ forces the degradation of mouse dihydrofolate reductase (DHFR), which is otherwise stable posttranslationally. We attached Rpl8A_1-50_ and Rpl8A_51-100_ to the N-terminus of mouse DHFR with a hemagglutinin (ha) epitope tag at the C-terminus for detection. Cycloheximide (CHX)-chase assay was performed to measure the stability of Rpl8A_1-50_-DHFRha and Rpl8A_51-100_-DHFRha, controlled by DHFRha and Rpl8A_1-100_-DHFRha. As shown in Fig. 1B, Rpl8A_1-50_-DHFRha was as stable as DHFRha during the chase. In contrast, Rpl8A_51-100_-DHFRha was rapidly degraded like Rpl8A_1-100_-DHFRha. At the beginning of chase (i.e., zero time point), the concentration of Rpl8A_51-100_-DHFRha was already much lower than that of DHFRha. Consistently, we found that the steady-state level of Rpl8A_51-100_-DHFRha was markedly lower than that of DHFRha (Fig. 1C, compare lanes 1, 4). The abundances of transcripts encoding DHFRha and the fusion proteins were comparable (Fig. S2A), indicating that the difference in protein levels is not caused by transcription. These experiments demonstrate that Rpl8A_51-100_, not Rpl8A_1-50_, carries the degron.

### The N-terminal IDR of Rpl8A is a stabilizer against CTPD

Contrary to the idea that IDRs are sensitive to proteolysis, Rpl8A_1-50_-DHFRha was not degraded during CHX chase. Instead, the steady-state level of Rpl8A_1-50_-DHFRha was much higher than that of DHFRha (Fig. 1C, compare lanes 1 and 3). We have shown that mouse DHFR, while posttranslationally stable, is degraded cotranslationally in yeast cells (30). Our results indicate that adding Rpl8A_1-50_ to the N-terminus inhibits CTPD of DHFR. We further compared the effect of Rpl8A_1-50_ with N-terminal Ub, which has been shown to inhibit CTPD of a variety of proteins including DHFR (25, 30). Yeast cells were transformed with plasmids expressing DHFRha, Rpl8A_1-50_-DHFRha, and Ub-DHFRha. As expected, yeast cells expressing the Ub-DHFRha fusion precursor produced much more DHFRha than the cells expressing DHFRha (Fig. 1D). (The N-terminal Ub is cleaved by Ub-specific proteases, releasing DHFRha.) The steady-state level of Rpl8A_1-50_-DHFRha was comparable to the DHFRha level in cells expressing Ub-DHFRha. Thus, N-terminal Rpl8A_1-50_ and Ub display a similar strength in suppressing CTPD of DHFRha.

We next examined if Rpl8A_1-50_ could inhibit CTPD of ribosomal protein Rps31, which is synthesized as a Ub fusion precursor (Ub-Rps31) in eukaryotic cells (31). We have shown that N-terminal Ub protects Rps31 from CTPD, as Rps31 made without N-terminal Ub is rapidly degraded cotranslationally (25). Consistent with early observations, the cells expressing Ub-Rps31ha produced much more Rps31ha than the cells expressing Rps31ha (Fig. 1E). The steady-state level of Rpl8A_1-50_-Rps31ha was also markedly higher than that of Rps31ha synthesized without N-terminal Ub. The transcript levels of *RPS31ha*, *UB-RPS31ha*, and *RPL8A_1-50_-RPS31ha* were comparable (Fig. S2B). These results demonstrate that N-terminal Rpl8A_1-50_ blocks CTPD of Rps31ha. We further assessed if deleting Rpl8A_1-50_ could enhance CTPD of Rpl8A. Indeed, the steady-state level of Rpl8A_Δ1-50_ha was lower than that of Rpl8Aha (Fig. 1F). Deletion of Rpl8A_1-50_ did not cause posttranslational degradation as Rpl8A_Δ1-50_ha was as stable as full-length Rpl8Aha during CHX chase (Fig. 1G). qRT-PCR analysis revealed no reduction in the abundance of *RPL8A_Δ1-50_ha* transcript compared to *RPL8Aha* (Fig. S2C). These results indicate that Rpl8A_1-50_ is a stabilizing element (stabilizer) impeding CTPD of Rpl8A. The Rpl8A_1-50_ stabilizer is portable, as it protects DHFR and Rps31 from CTPD when placed to their N-termini.

We wondered if the stabilizer of Rpl8A would also increase the steady-state levels of proteins that are not degraded cotranslationally. To address this question, we evaluated if adding Rpl8A_1-50_ to the N-terminus of Rps12 would increase its expression level. Our previous study showed that Rps12 is not subject to CTPD when it is expressed at a moderate level from the *CUP1* promoter (25). Consistent with our early results, yeast cells expressing Ub-Rps12ha produced a similar amount of Rps12ha as the counterparts expressing Rps12ha (Fig. 1H, compare lanes 1, 2), confirming no CTPD of Rps12ha. The steady-state level of Rpl8A_1-50_-Rps12ha was also comparable to that of Rps12ha (compare lanes 1 and 3), indicating that N-terminal Rpl8A_1-50_ does not increase the expression of Rps12ha. The green fluorescent protein (GFP) is another protein that is not subject to CTPD, as yeast cells expressing Ub-GFP fusion precursor yielded a similar amount of GFP as the cells expressing GFP (Fig. 1I, compare lanes 1 and 2). The addition of Rpl8A_1-50_ to the N-terminus of GFP also did not increase the steady-state level (compare lanes 2 and 3). Thus, N-terminal Rpl8A_1-50_ has no impact on the expression of proteins that are not degraded cotranslationally. These results also indicate that adding Rpl8A_1-50_ to the N-terminus *per se* does not enhance translation speed or efficiency.

### The stabilizer of Rpl8A has no impact on posttranslational degradation

We showed that Rpl8A_1-100_-DHFRha was rapidly degraded during CHX-chase (Fig. 1B). This suggests that Rpl8A_1-50_ does not impede Rpl8A_51-100_-mediated posttranslational degradation of DHFR. We wanted to examine if the stabilizer of Rpl8A would inhibit posttranslational degradation directed by another degron. Our early work showed that residues 172-229 of yeast transcription factor Rpn4 (Rpn4_172-229_) constitute a portable degron for posttranslational degradation (32). We constructed three plasmids to express Rpn4_172-229_-DHFRha, Rpl8A_1-50_-Rpn4_172-229_-DHFRha, and Ub-Rpn4_172-229_-DHFRha, respectively (Fig. 2A). The abundances of transcripts generated from these 3 plasmids were comparable (Fig. 2B). CHX-chase analysis was performed to measure posttranslational degradation of these 3 fusion proteins (Fig. 2C). Consistent with our previous observations, Rpn4_172-229_-DHFRha was unstable during the 30 min chase period (lanes 1, 2). Rpl8A_1-50_-Rpn4_172-229_-DHFRha was also rapidly degraded (lanes 3, 4). Therefore, Rpl8A_1-50_ does not interfere with posttranslational degradation driven by the Rpn4_172-229_ degron. Notably, the level of Rpl8A_1-50_-Rpn4_172-229_-DHFRha was higher than that of Rpn4_172-229_-DHFRha at the beginning of chase (compare lanes 1, 3). The steady-state level of Rpl8A_1-50_-Rpn4_172-229_-DHFRha was indeed higher than that of Rpn4_172-229_-DHFRha (Fig. 2D, compare lanes 2, 3). These results indicate that Rpn4_172-229_-DHFRha is also subject to CTPD, which is inhibited by N-terminal Rpl8A_1-50_. Similarly, adding Ub to the N-terminus had no effect on posttranslational degradation of Rpn4_172-229_-DHFRha (Fig. 2C, lanes 5, 6), but blocked its CTPD (Fig. 2D, compare lanes 2, 4). These experiments demonstrate that the stabilizer of Rpl8A has no impact on posttranslational degradation.

**Figure 2.**
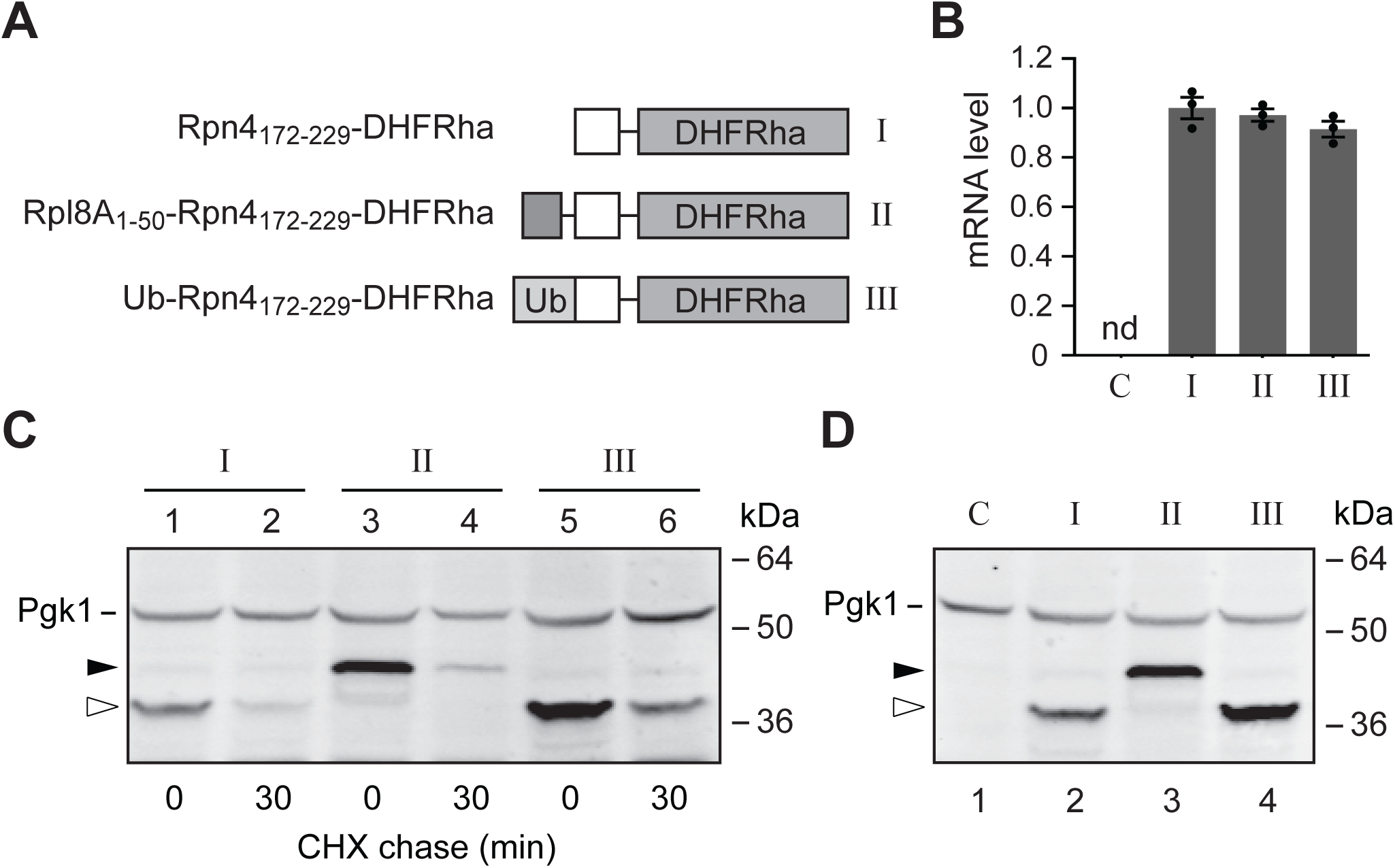
The stabilizer of Rpl8A has no impact on posttranslational protein degradation. (A) Schematic of fusion proteins Rpn4_172-229_-DHFRha (I), Rpl8A_1-50_-Rpn4_172-229_-DHFRha (II), and Ub-Rpn4_172-229_-DHFRha (III). All proteins were expressed under the control of *CUP1* promoter from a low-copy number vector. (B) qRT-PCR analysis of the transcripts encoding fusion proteins I, II and III. A yeast transformant with an empty vector served as a negative control. Primers YX1084 and YX1018 were used for PCR. Data are presented as mean ± S.D. from 3 independent experiments. (C) Rpl8A_1-50_ has no effect on posttranslational degradation. CHX chase analysis was performed for yeast transformants expressing proteins I-III. Rpn4_172-229_-DHFRha and Rpl8A_1-50_-Rpn4_172-229_-DHFRha were marked by open and filled arrowheads, respectively. (D) Rpl8A_1-50_ suppresses CTPD of Rpn4_172-229_-DHFRha. Western blotting was performed to compare the steady-state levels of Rpn4_172-229_-DHFRha (lanes 2, 4) and Rpl8A_1-50_-Rpn4_172-229_-DHFRha (lane 3). The yeast transformant with a void vector was used as a negative control (lane 1).

### Rpl8A_1-50_ is bound by HSP70 chaperone proteins Ssa and Ssb

We wanted to elucidate the mechanism by which Rpl8A_1-50_ protects proteins from CTPD. Unlike N-terminal Ub, which folds cotranslationally and probably blocks the access of proteasomes to nascent chains, Rpl8A_1-50_ is a floppy fragment. How can an unstructured fragment, which is considered a preferred target for proteasomal degradation, function as a stabilizer against CTPD? To address this question, we decided to identify proteins bound to Rpl8A_1-50_ using a pulldown-mass spectrometry (LC-MS/MS) approach. We constructed a plasmid to produce Rpl8A_1-50_-GSTFlag, a fusion protein with Rpl8A_1-50_ attached to the N-terminus of glutathione S-transferase (GST) carrying a C-terminal Flag tag. This plasmid and a control vector expressing GSTFlag were transformed into yeast cells. The steady-state level of Rpl8A_1-50_-GSTFlag was much higher than that of GSTFlag (Fig. 3A). CHX-chase analysis showed that both GSTFlag and Rpl8A_1-50_-GSTFlag were posttranslationally stable (Fig. 3B). These results indicate that GSTFlag is subject to CTPD, which is inhibited by N-terminal Rpl8A_1-50_. Cell extracts from yeast transformants expressing GSTFlag and Rpl8A_1-50_-GSTFlag were passed through glutathione agarose beads. Retained proteins were resolved by SDS-PAGE and stained with Coomassie blue (Fig. 3C). Note that more GST-Flag lysate was applied in the pulldown to ensure that comparable amounts of GSTFlag and Rpl8A_1-50_-GSTFlag were loaded on the beads. Two predominant bands (*a* and *b*) were obtained from the pulldown of Rpl8A_1-50_-GSTFlag but not GSTFlag. LC-MS/MS analysis revealed that band *a* included Ssa1, Ssa2, and Ssa4 proteins, whereas band *b* contained Ssb1 and Ssb2 proteins (Fig. S3). Ssa1, Ssa2, and Ssa4 are 3 of the 4 SSA subfamily members of HSP70 chaperone proteins (33). Ssa3 is not expressed under normal growth conditions, thus its absence is not noteworthy. Ssa1 and Ssa2 are paralogs with 98% identity and share 84% identity with Ssa4. The paralogs Ssb1 and Ssb2 are 99% identical and form the SSB subfamily of HSP70 (34).

**Figure 3.**
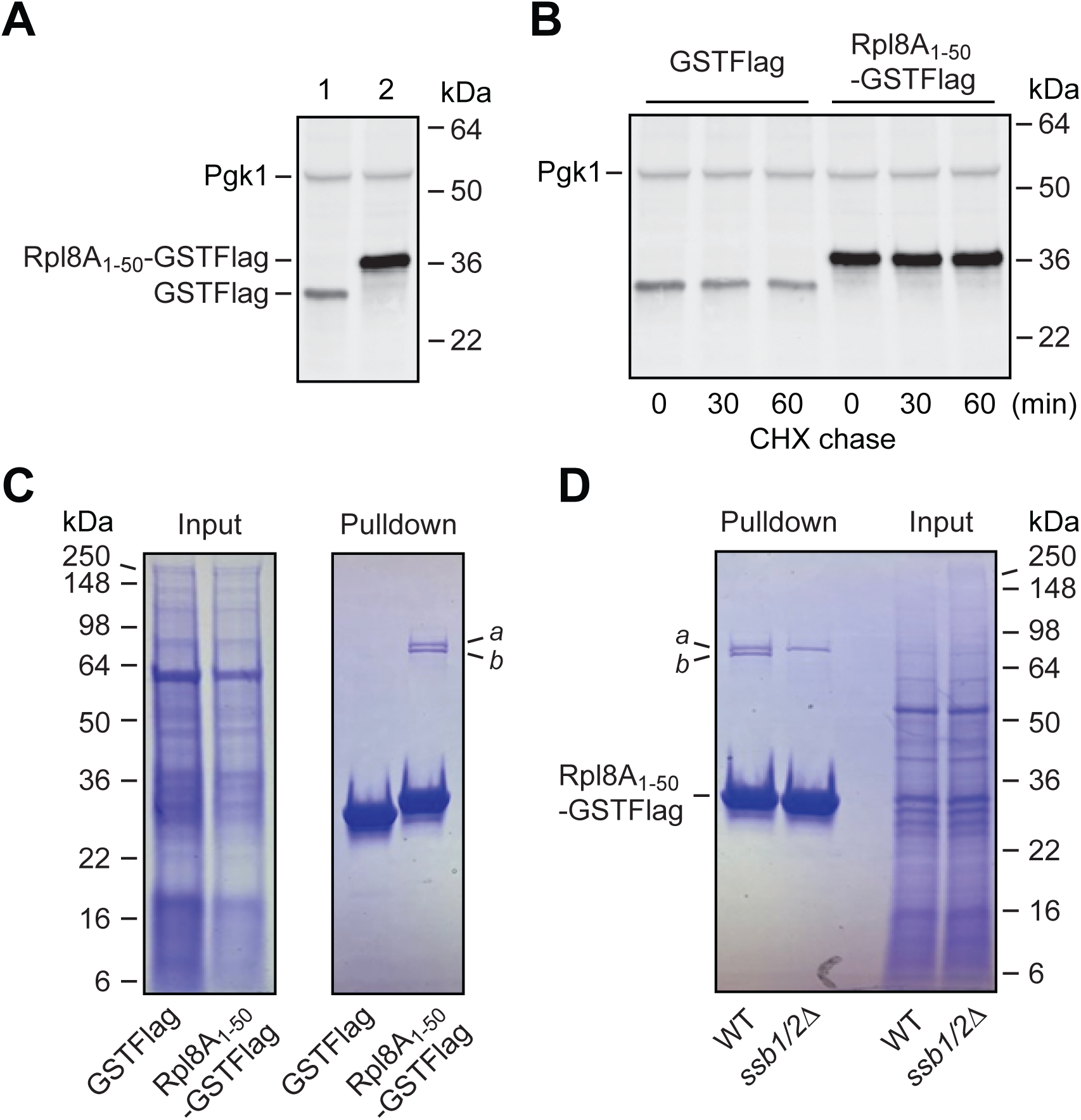
Rpl8A_1-50_ is bound by chaperone proteins Ssa and Ssb. (A) Comparison of the steady-state levels of GSTFlag and Rpl8A_1-50_-GSTFlag. Yeast cells expressing GSTFlag and Rpl8A_1-50_-GSTFlag were analyzed by Western blotting with anti-Flag and anti-Pgk1 antibodies. (B) GSTFlag and Rpl8A_1-50_-GSTFlag are posttranslationally stable. CHX chase analysis was conducted to monitor posttranslational degradation of GSTFlag and Rpl8A_1-50_-GSTFlag. (C) Identification of proteins bound to Rpl8A_1-50_-GSTFlag. Cell extracts from yeast transformants expressing GSTFlag or Rpl8A_1-50_-GSTFlag were applied to GST pulldown assay. More GSTFlag lysate was used to ensure similar amounts of GSTFlag and Rpl8A_1-50_-GSTFlag in the input (left panel). Retained proteins were separated by SDS-PAGE, followed by Coomassie Blue staining (right panel). Bands *a* and *b* specifically pulled down by Rpl8A_1-50_-GSTFlag were sliced and subjected to protein identification by LC-MS/MS analysis (see Fig. S3). (D) Ssb1/Ssb2 proteins specifically pulled down by Rpl8A_1-50_-GSTFlag. Extracts from WT and *ssb1/2Δ* cells expressing Rpl8A_1-50_-GSTFlag were applied to pulldown assay. Retained proteins were detected as C. Band *b* was absent from the pulldown of *ssb1/2Δ* mutant.

To validate the LC-MS/MS results, we transformed the Rpl8A_1-50_-GSTFlag plasmid into a mutant strain (*ssb1/2Δ*) deleted of both *SSB1* and *SSB2* genes and repeated the pulldown assay. A WT strain expressing Rpl8A_1-50_-GSTFlag was used as a control. As shown in Fig. 3D, band *b* was absent from the pulldown of the *ssb1/2Δ* mutant. This experiment confirms the binding of Ssb1/2 proteins to Rpl8A_1-50_. There is no viable yeast strain with *SSA1*, *SSA2* and *SSA4* genes deleted simultaneously (32). Therefore, it is not possible to perform a similar pulldown assay using a null mutant lacking Ssa1, Ssa2, and Ssa4 proteins. The identification of Ssa/Ssb bound to Rpl8A_1-50_ is interesting because they are known to assist cotranslational folding of nascent polypeptides (33–42). Our results suggest that Rpl8A_1-50_ may suppress CTPD through recruiting HSP70 chaperone proteins to nascent chains, thereby facilitating cotranslational folding.

### Suppression of CTPD by other N-terminal IDRs

We next examined if other N-terminal IDRs could also act as a stabilizer against CTPD. To this end, we took advantage of the AlphaFold Protein Structure Database integrated into the Saccharomyces Genome Database to acquire predicted 3D structures of yeast proteins. For this study, we tested 9 proteins (Ape2, Spn1, Ero1, Put1, Bgl2, Sso2, Hrk1, Mrps9, and Rck2), all having a significant N-terminal IDR (Fig. 4A). Our previous study showed that these proteins are not or only marginally degraded cotranslationally (23). Because HSP70 typically bind to the core unfolded peptide sequence of 5 to 7 amino acids (43), we decided to attach the first 10 amino acids of the proteins to be tested to the N-terminus of DHFRha. Of note, these 10-residue IDRs do not share noticeable homology of amino acid sequences (Fig. S4). N-terminal Ub was used as a reference in this experiment. We found that all 9 IDRs suppressed CTPD of DHFRha (Fig. 4B). The steady-state levels of the IDR-DHFRha fusion proteins were 1.7-3.5 times as high as that of DHFRha (Fig. 4C). Some of the IDRs displayed a comparable strength as the N-terminal Ub. These results suggest that suppression of CTPD is likely a generic function of N-terminal IDRs.

**Figure 4.**
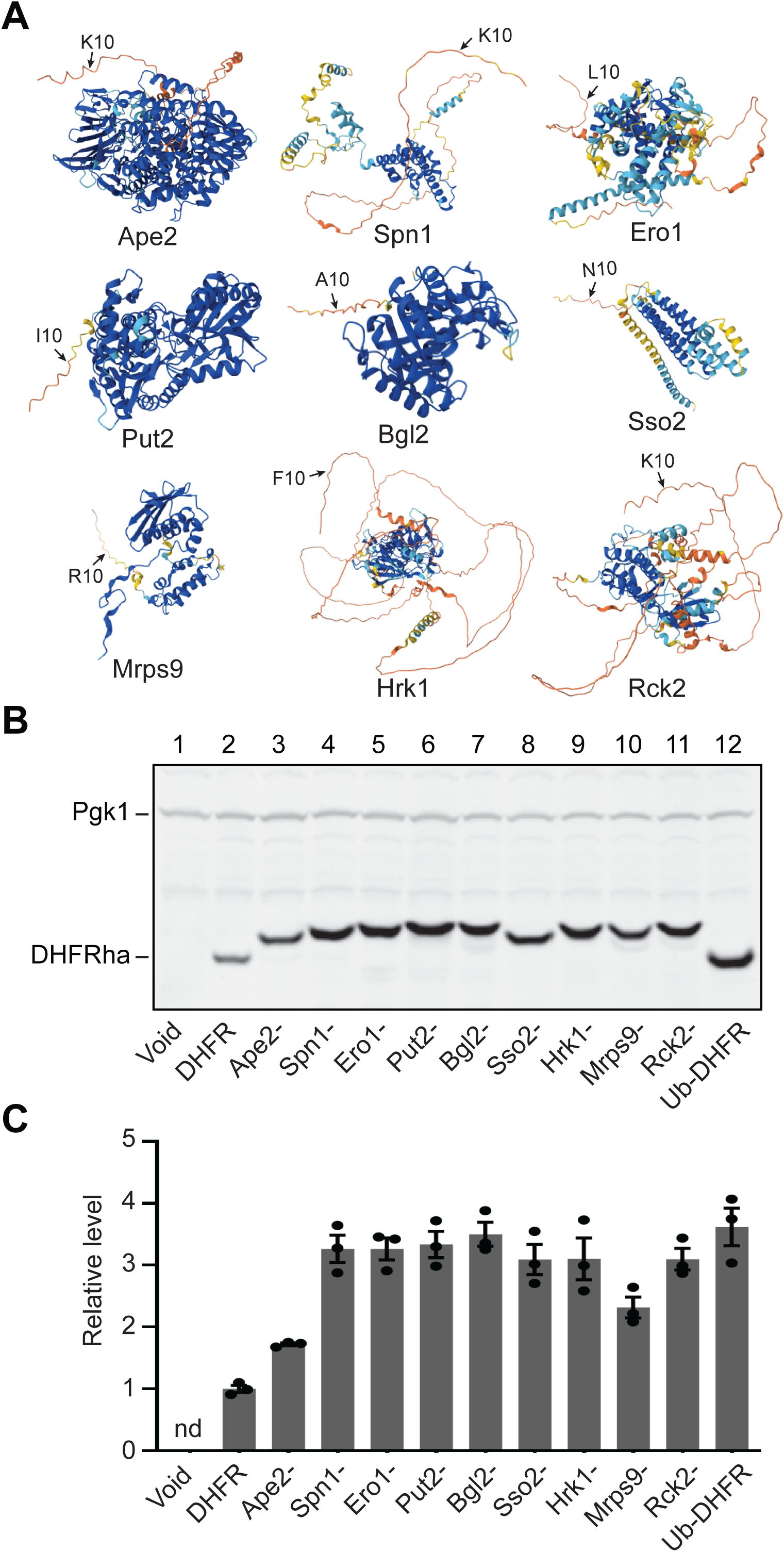
Suppression of CTPD by different N-terminal IDRs. (A) Predicted protein structures. The predicted structures of 9 yeast proteins were obtained from the AlphaFold Protein Structure Database integrated into the Saccharomyces Genome Database (https://www.yeastgenome.org). The tenth residue of each protein was marked. (B) Comparison of the steady-state levels of DHFRha and fusion proteins carrying various N-terminal 10-residue IDRs as indicated. Western blotting analysis was performed with anti-ha and anti-Pgk1 antibodies. (C) Quantification of Western blots. Western blots were quantified by Image J. The levels of DHFRha and IDR-DHFRha fusion proteins were normalized against Pgk1. Shown are the relative ratios of IDR-DHFRha to DHFRha. Data are presented as mean ± S.D. from 3 independent experiments.

### The N-terminal IDR of hRpl7A suppresses CTPD of CFTR in human cells

We wanted to examine if N-terminal IDR-mediated suppression of CTPD also occurs in human cells. The human ribosomal protein homologous to yeast Rpl8A is hRpl7A (44). hRpl7A shares 55% identity with Rpl8A in amino acid sequence (Fig. S5). Structural analysis by AlphaFold revealed that the N-terminal region of hRpl7A including the first 55 amino acids (hRpl7A_1-55_) is an IDR (Fig. 5A). We expressed C-terminally V5-tagged hRpl7A and a truncation mutant (hRpl7A_Δ1-54_) deleted of the first 54 amino acids in human embryonic kidney 293 (HEK293) cells. While hRpl7AV5 was easily detected by Western blotting, hRpl7A_Δ1-54_V5 was virtually undetectable (Fig. S6A). The transcript encoding hRpl7A_Δ1-54_V5 was more abundant than the hRpl7AV5 counterpart (Fig. S6B). These results indicate that the N-terminal IDR protects hRpl7A from CTPD.

**Figure 5.**
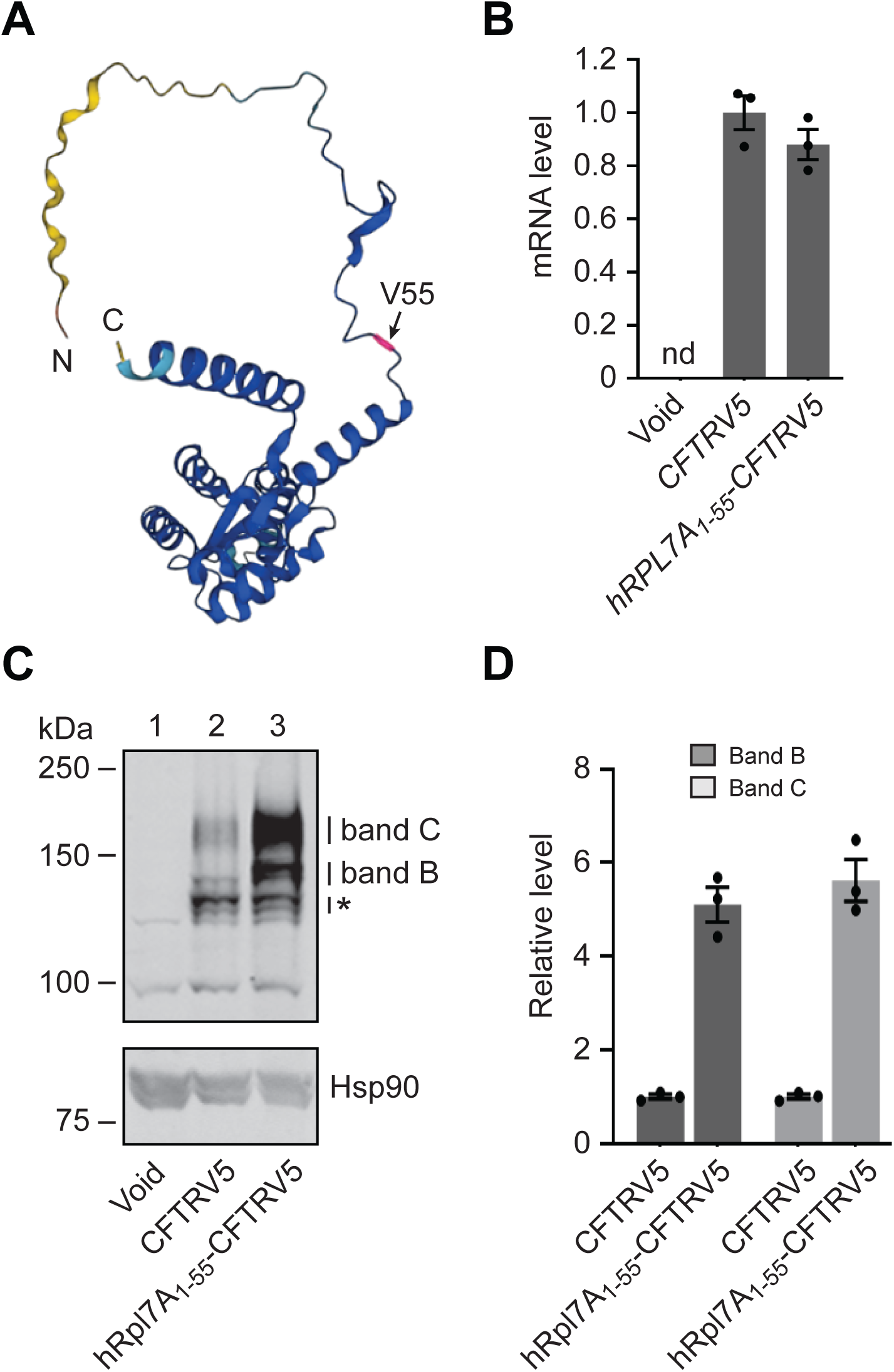
The N-terminal IDR of hRpl7A protects CFTR from CTPD in human cells. (A) Structure analysis of hRpl7A by AlphaFold. The structure of hRpl7A was obtained from Protein Data Bank (P62424). Val-55 was marked by an arrow. (B) qRT-PCR analysis of the transcripts encoding CFTRV5 and hRpl7A_1-55_-CFTRV5. HEK293 cells with mock transfection served as a negative control. PCR was performed with primers YX1104 and YX1105 corresponding to the V5 tag sequence and the BGH reverse priming site on vector pEF6/V5-His A. U6 was used as an internal control for normalization of gene expression. Data are presented as mean ± S.D. from 3 independent experiments. (C) N-terminal hRpl7A_1-55_ increased the abundances of band B and band C of CFTR. Cells expressing CFTRV5 (lane 2), hRpl7A_1-55_-CFTRV5 (lane 3), or a void vector (lane 1) were analyzed by Western blotting with anti-V5 and anti-Hsp90 antibodies. Hsp90 served as an internal control. B and C bands were indicated. The asterisk marked cleavage products of CFTR. (D) Quantification of Western blots. Western blots were quantified by Image J. The levels of B and C bands of CFTRV5 and hRpl7A_1-55_-CFTRV5 were normalized against Hsp90. Shown are the relative abundances of hRpl7A_1-55_-CFTRV5 versus CFTRV5. Data are presented as mean ± S.D. from 3 independent experiments.

We next tested if hRpl7A_1-55_ could act as a stabilizer against CTPD of CFTR, a well-known substrate for CTPD (45, 46). During synthesis on the endoplasmic reticulum (ER)-associated ribosomes, CFTR is translocated into the ER and undergoes cotranslational folding and core glycosylation at two asparagine residues (N894 and N900) (47). Folded, core-glycosylated CFTR is then trafficked through the secretory pathway to the Golgi apparatus where it is further glycosylated and matured. The folding of nascent CFTR is extremely inefficient and subject to extensive quality control. As a result, only a small fraction of CFTR completes folding and is exported to Golgi, whereas the majority of CFTR is degraded prematurely in the ER by the Ub-proteasome system (48–50). We transfected HEK293 cells with plasmids encoding C-terminally V5-tagged CFTR and hRpl7A_1-55_-CFTR. The transcript level of hRpl7A_1-55_-CFTRV5 was relatively lower than that of CFTRV5 (Fig. 5B). The characteristic B and C bands of CFTR, representing core glycosylated immature form and complex glycosylated mature form, were detected in both transfectants by Western blotting (Fig. 5C, Fig. S7). This demonstrates that the N-terminal hRpl7A_1-55_ does not interfere with the processing of CFTR such as glycosylation and trafficking. Remarkably, the levels of band B and band C of hRpl7A_1-55_-CFTRV5 were at least five times as high as the CFTRV5 counterparts (Fig. 5D). Thus, adding hRpl7A_1-55_ to the N-terminus of CFTR reduces its CTPD by more than 80%.

## Discussion

Efficient folding initiated from the N-terminal region is conceivably critical for nascent chains to resist CTPD because a folded N-terminal region may obstruct the access of proteasome. An excellent example is the N-terminal Ub moiety that protects Ub-fusion proteins from CTPD (25). On the other hand, nascent polypeptides with an N-terminal IDR are thought to be susceptible to CTPD. Contrary to this assumption, we found that N-terminal IDRs suppress CTPD. This function may be generic for N-terminal IDRs because the N-terminal IDRs of all 10 yeast proteins tested in this study inhibit CTPD. Moreover, the N-terminal IDR of human ribosomal protein hRpl7A also acts as a stabilizer against CTPD in human cells, indicating that suppression of CTPD by N-terminal IDRs is conserved from yeast to human.

N-terminal IDRs are unable to fold and therefore cannot block the access of proteasomes to nascent chains by themselves. As such, they must use a different mechanism from N-terminal Ub in suppressing CTPD. We demonstrated that Rpl8A_1-50_-GSTFlag but not GSTFlag is bound by Ssa/Ssb proteins and that Rpl8A_1-50_-GSTFlag is resistant whereas GSTFlag is sensitive to CTPD (Fig. 3). We also showed that adding hRpl7A_1-55_ to the N-terminus of CFTR drastically reduces its CTPD. It is well documented that CTPD of CFTR results from extremely inefficient cotranslational folding (45–50). These observations suggest that Rpl8A_1-50_, hRpl7A_1-55_ and other IDRs may suppress CTPD through recruiting HSP70 chaperone proteins to nascent chains, which are known to assist nascent chain folding (33–42).

The finding that N-terminal IDRs inhibit CTPD raises an interesting question: why a protein with a foldable N-terminal region is less resistant to CTPD than one carrying an N-terminal IDR? For example, AlphaFold reveals that the N-terminal region of DHFR forms a helix. Yet, adding an IDR to its N-terminus reduces CTPD. Similarly, the Rpl8A_Δ1-50_ mutant missing the N-terminal IDR has a helical structure at the N-terminus, but it is subject to higher level CTPD than Rpl8A. Although further investigation is required to explicitly address the underlying mechanism, there are several possible explanations. First, not all “foldable” N-terminal regions can cotranslationally fold as efficiently and completely as N-terminal Ub. A partially folded N-terminal domain may not be sufficient to block the access of proteasome, and the nascent polypeptide chain is still subject to CTPD. Second, a partially folded N-terminal region likely has lower affinity for HSP70 than an IDR. It is well known that HSP70 chaperone proteins preferably bind to unfolded nascent chains (36, 37). Therefore, partial folding at the N-terminus may actually reduce the chance for an elongating nascent chain to acquire and/or retain chaperone proteins to secure its complete folding. Third, in addition to assisting cotranslational folding, the chaperone proteins recruited by N-terminal IDRs may block proteasomes from attacking nascent chains.

The finding of the role of N-terminal IDRs in suppressing CTPD has potential translational impacts. Using an N-terminal IDR, one can increase the yield of mature proteins such as CFTR that are otherwise difficult to produce due to inefficient cotranslational folding and high level CTPD. It may also be possible to develop IDR-based therapeutics for diseases caused by inappropriate protein folding.

## Methods

### Plasmids, yeast strains, human cells, and antibodies

The plasmids used in this study are listed in Table S1. All plasmids were verified by restriction enzyme digestion and DNA sequencing. For yeast expression vectors, target proteins were expressed from the copper-inducible *CUP1* promoter in vectors pRS314 or pRS426. For mammalian expression vectors, target proteins were expressed under the human elongation factor 1α-subunit promoter from the vector PEF6/V5-His A (Invitrogen). Construction of yeast expression vectors for Ub-fusion precursor proteins was described previously (25). The yeast strains used in this study include JD52 (*MAT***a** *trp1-Δ63 ura3-52 his3-Δ200 leu2-3,112 lys2-801*), DS10 (*MAT***a** *lys1 lys2 leu2-3*,*112 ura3-52 his3-11*,*15 trp1-*Δ*1*), *ssa1/2Δ* (*ssa1Δ*::*HIS3 ssa2Δ*::*LEU2* derivative of DS10), and *ssb1/2Δ* (*ssb1Δ*::*HIS3 ssb2Δ*::*LEU2* derivative of DS10). Strains DS10, *ssa1/2Δ* and *ssb1/2Δ* were generated by Elizabeth Craig’s laboratory at University of Wisconsin (32, 33). JD52 was described previously (51). Human embryonic kidney 293 (HEK293) cells from the American Type Culture Collection authenticated by short tandem repeat profiling were used to express mammalian proteins. The anti-Flag and anti-β-actin monoclonal antibodies were obtained from Sigma-Aldrich. The anti-GFP and anti-V5 monoclonal antibodies were produced by Invitrogen. The anti-ha monoclonal, anti-Pgk1 rabbit polyclonal, and anti-heat shock protein 90 (Hsp90) rabbit polyclonal antibodies were purchased from Covance, OriGene, and ProteinTech, respectively. Antibodies were used according to the manufacturers’ instructions.

### CHX-chase assay and Western blotting analysis

Yeast cells carrying *CUP1* promoter-based plasmids were cultured in synthetic defined medium supplemented with essential amino acids. Exponentially growing cells were diluted to OD_600_ of ∼0.2 and induced with CuSO_4_ (0.1 mM) for 8 h before harvested for Western blotting analysis. For CHX-chase assay, CHX was added to cell cultures grown to OD_600_of ∼ 0.8 at a final concentration of 0.2 mg/ml. Equal volumes of samples were withdrawn at each time point during the chase. Cells were lysed by incubation with 2x SDS buffer (2% SDS, 30 mM dithiothreitol, 90 mM Na-HEPES, pH 7.5) at 100°C for 5 min. Supernatants were recovered by centrifugation and subjected to Western blotting as described previously (30). The blots were probed simultaneously with mouse monoclonal antibodies against epitope-tagged target proteins and the anti-Pgk1 rabbit polyclonal antibody. The house-keeping enzyme phosphoglycerate kinase (Pgk1) served as a loading control. After removal of unbound antibodies, the blots were incubated with fluorescent dye (Alexa Fluor 680)-conjugated goat-anti-mouse and donkey-anti-rabbit secondary antibodies, followed by detection with the Odyssey infrared imaging system (Li-Cor Biosciences). For human cell experiments, HEK293 cells transfected with various plasmids were harvested 28 h after transfection for Western blotting analysis. HEK293 transfectants were lysed with 2x SDS buffer. Chromosomal DNA was sheared to lower viscosity by passing the lysates through 28-gauge needles. Blots were first probed with anti-V5 antibody and Alexa Fluor 680-conjugated secondary antibody and subsequently re-probed with anti-Hsp90 antibody, followed by Alexa Fluor 800-conjugated secondary antibody.

### GST pulldown assay and protein identification by mass spectrometry

Yeast cells expressing GSTFlag or Rpl8A_1-50_-GSTFlag from the *CUP1* promoter were induced by CuSO_4_ as described above. Harvested cells were resuspended in pulldown buffer (150 mM NaCl, 50 mM HEPES, pH 7.5, 10% glycerol, 0.1% Triton X-100) plus protease inhibitor mix (Roche Diagnostics) and broken by the SuperFastPrep-2 homogenizer. Supernatants were collected after centrifugation at 20,000 rpm for 10 min. Approximately 0.5 mg of cell lysate containing GSTFlag and ∼ 0.2 mg of cell lysate containing Rpl8A_1-50_-GSTFlag were incubated with glutathione-agarose beads at 4°C for 2 h. The beads were washed three times with pulldown buffer. The retained proteins were resolved by SDS-PAGE and stained with Coomassie Blue. Proteins specifically bound to Rpl8A_1-50_-GSTFlag were cut out. The gel slices were subjected to LC-MS/MS analysis for protein identification, following the same procedure as described previously (52).

### Quantitative reverse-transcription PCR (qRT-PCR)

The procedure of RNA isolation from yeast cells was described previously (25). RNeasy Kit (Qiagen) was used to prepare RNA from HEK293 transfectants. Reverse-transcription was conducted using SuperScript III reverse transcriptase (Invitrogen), and qRT-PCR was conducted with Fast SYBR Green Master Mix in the StepOne™ system according to the manufacturer’s instruction (Applied Biosystems). Experiments were performed in triplicate. The *ACT1* and U6 housekeeping genes served as internal controls for normalization of gene expression levels in yeast and HEK293 cells, respectively. The oligo primers for qRT-PCR are listed in Table S2.

### Deglycosylation of CFTR

Glycosidases Endo H and PNGase F were purchased from New England Biolabs. Cell lysates were treated with Endo H and PNGase F according to the manufacturer’s instructions. After treatments, proteins were precipitated with 4 fold cold ethanol and centrifugation at 20,000 rpm for 10 min. The pellets were washed with 80% ethanol and dissolved with 1x sample buffer. Protein samples were separated by SDS-PAGE, followed by Western blotting.

## Statistical analysis

Statistical analyses were performed using one-way analysis of variance (ANOVA). All data were presented as mean ± standard deviation (S.D.) from three independent experiments.

## Data Availability

The data supporting the findings of this study are in the manuscript and the Supplementary Information appendix.

## Acknowledgments

We thank Elizabeth Craig, Kevin Morano, Fei Sun, Junying Zhou and Jing Li for yeast strains, antibodies, plasmids and cell lines. We also thank Kang Chen for comments on the manuscript.

## Author contributions

DJ, DX, and YX developed the conceptual framework and designed experiments; DJ, DX, JW, SW and YX performed experiments; DJ, DX, JW, SW, LL and YX analyzed data; DJ, DX and YX wrote the original manuscript; DJ, DX, JW, SW, LL and YX reviewed and edited the manuscript.

## Funding and additional information

This work was supported by Karmanos Cancer Institute at Wayne State University to YX.

