## Supporting Information for "Intrinsically disordered N-terminal regions suppress cotranslational protein degradation"

**This PDF file includes:**

Figures S1 to S7
Tables S1 to S2

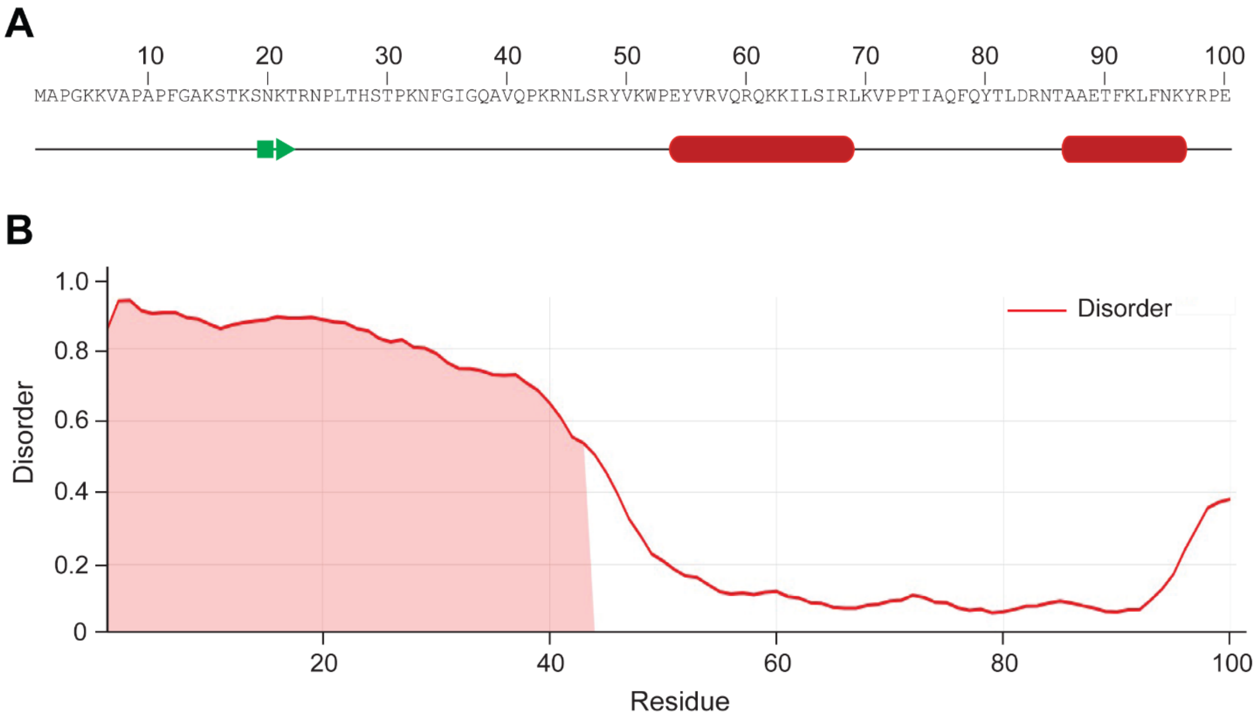


**Figure S1. The N-terminal region of Rpl8A is unstructured.** (A) Predicted secondary structure of the Rpl8A$_{1\text{-}100}$ fragment. Prediction was conducted by the JPrep server. The green arrow marks a potential short sheet, whereas the red oblongs indicate helices. (B) Plots of per-residue disorder for Rpl8A$_{1\text{-}100}$ by Metapredict V2. Shown are the AlphaFold structural confidence scores (between 0 to 1) for each residue. A score higher than 0.5 means that more than half of the predictors predict the residue to be disordered.

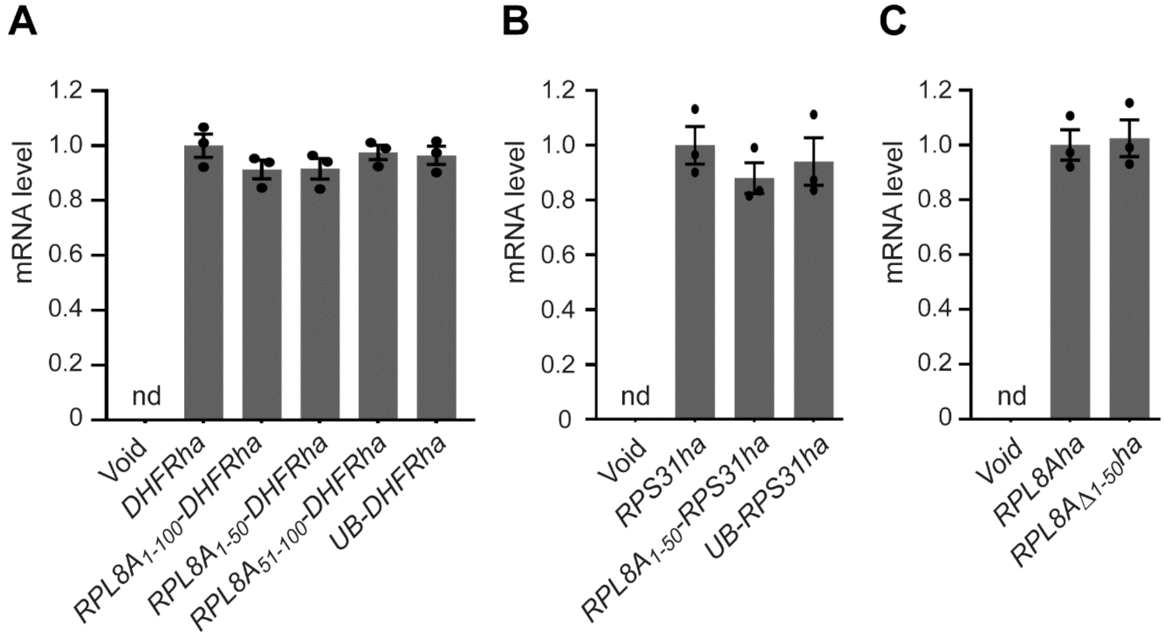


**Figure S2. qRT-PCR analyses.** (A) Comparable abundances of transcripts encoding DHFRha, Rpl8A$_{1-100}$-DHFRha, Rpl8A$_{1-50}$-DHFRha, Rpl8A$_{51-100}$-DHFRha, and Ub-DHFRha. Primers YX1084 and YX1018 were used for PCR. YX1084 is ~ 100 bp upstream from the stop codon of *DHFR*, whereas YX1018 corresponds to the C-terminal ha sequence. The yeast transformant carrying a void vector (pRS314) served as a negative control. (B) Measurement of the abundances of transcripts encoding Rps31ha, Rpl8A$_{1-50}$-Rps31ha, and Ub-Rps31ha. Primers YX1014 and YX1018 were used for PCR. YX1014 is ~ 175 bp upstream from the stop codon of *RPS31*. (C) Comparison of the levels of transcripts encoding Rpl8Aha and Rpl8A$_{\Delta 1-50}$ha. Primers YX1034 and YX1018 were used for PCR. YX1034 is ~170 bp upstream from the stop codon of *RPL8A*. The *ACT1* housekeeping gene served as an internal control for normalization of gene expression levels. All data are presented as mean ± S.D. from three independent experiments.

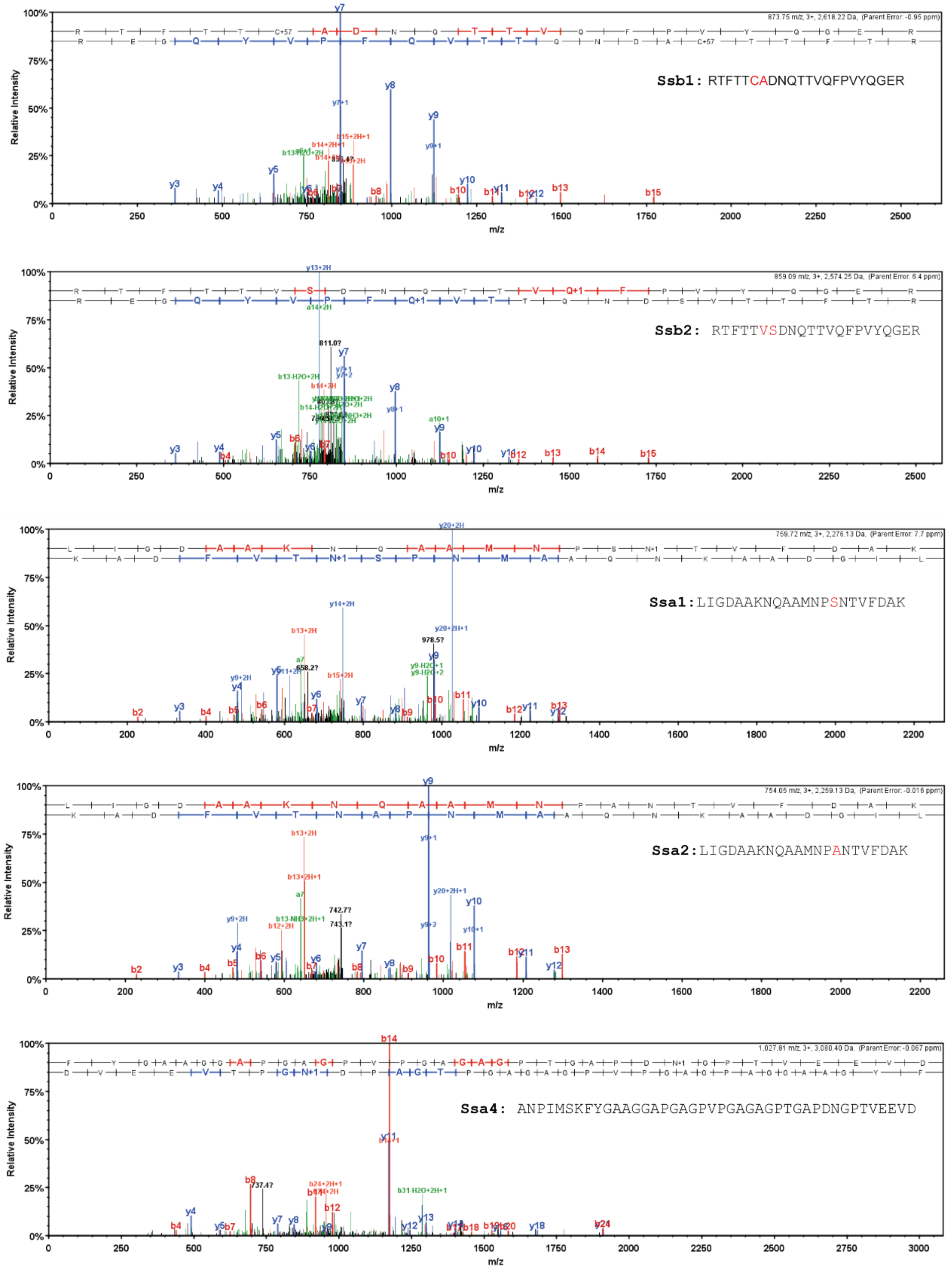


**Figure S3. Identification of Ssa and Ssb proteins by LC-MS/MS.** The bands *a* and *b* pulled down specifically by Rpl8A$_{1\text{-}50}$-GST-Flag (see Fig. 4C) were analyzed by LC-MS/MS. Shown here are the representative spectra of peptides derived from the top five proteins with the highest hits, including Ssb1, Ssb2, Ssa1, Ssa2, and Ssa4. The residues distinguishing Ssb1 from Ssb2, and Ssa1 from Ssa2 are labelled in red.

```
        1        10
Ape2:   MPIVRWLLLK
Spn1:   MSTADQEQPK
Ero1:   MRLRTAIATL
Put2:   MLSARCLKSI
Bgl2:   MRFSTTLATA
Sso2:   MSNANPYENN
Hrk1:   MPNLLSRNPF
Mrps9:  MFSRLSLFRR
Rck2:   MLKIKALFSK
```

**Figure S4. Alignment of the N-terminal 10 amino acids of 9 proteins as indicated.**

```
  Yeast Rpl8A  1     M-----APGKKVAPAPFGAKSTKSNKTRNPLTHSTPKNFGIGQAVQPKRNLSRYVKWPEY  55
                     M     A GKKVAPAP   K  ++ K  NPL    PKNFGIGQ +QPKR+L+R+VKWP Y
  Human Rpl7A  1     MPKGKKAKGKKVAPAPAVVKKQEAKKVVNPLFEKRPKNFGIGQDIQPKRDLTRFVKWPRY  60

  Yeast Rpl8A  56    VRVQRQKKILSIRLKVPPTIAQFQYTLDRNTAAETFKLFNKYRPETAAEKKERLTKEAAA  115
                     +R+QRQ+ IL  RLKVPP I QF   LDR TA +  KL +KYRPET  EKK+RL   A
  Human Rpl7A  61    IRLQRQRAILYKRLKVPPAINQFTQALDRQTATQLLKLAHKYRPETKQEKKQRLLARAEK  120

  Yeast Rpl8A  116   VAEGKSKQDASPKPYAVKYGLNHVVALIENKKAKLVLIANDVDPIELVVFLPALCKKMGV  175
                      A GK       +P  ++ G+N V  L+ENKKA+LV+IA+DVDPIELVVFLPALC+KMGV
  Human Rpl7A  121   KAAGKGDVPTK-RPPVLRAGVNTVTTLVENKKAQLVVIAHDVDPIELVVFLPALCRKMGV  179

  Yeast Rpl8A  176   PYAIVKGKARLGTLVNQKTSAVAALTEVRAEDEAALAKLVSTIDANFADKYDEVKKHWGG  235
                     PY I+KGKARLG LV++KT    A T+V +ED+ ALAKLV  I  N+ D+YDE+++HWGG
  Human Rpl7A  180   PYCIIKGKARLGRLVHRKTCTTVAFTQVNSEDKGALAKLVEAIRTNYNDRYDEIRRHWGG  239

  Yeast Rpl8A  236   GILGNKAQAKMDK--RAKNSDSA----  256
                      +LG K+ A++ K  +AK  + A
  Human Rpl7A  240   NVLGPKSVARIAKLEKAKAKELATKLG  266
```

**Figure S5. Sequence alignment between yeast Rpl8A and human hRpl7A.** The BLAST program of National Center for Biotechnology Information (NCBI) was used to align the protein sequences of Rpl8A and hRpl7A. These two sequences share 55% identities and 69% positives.

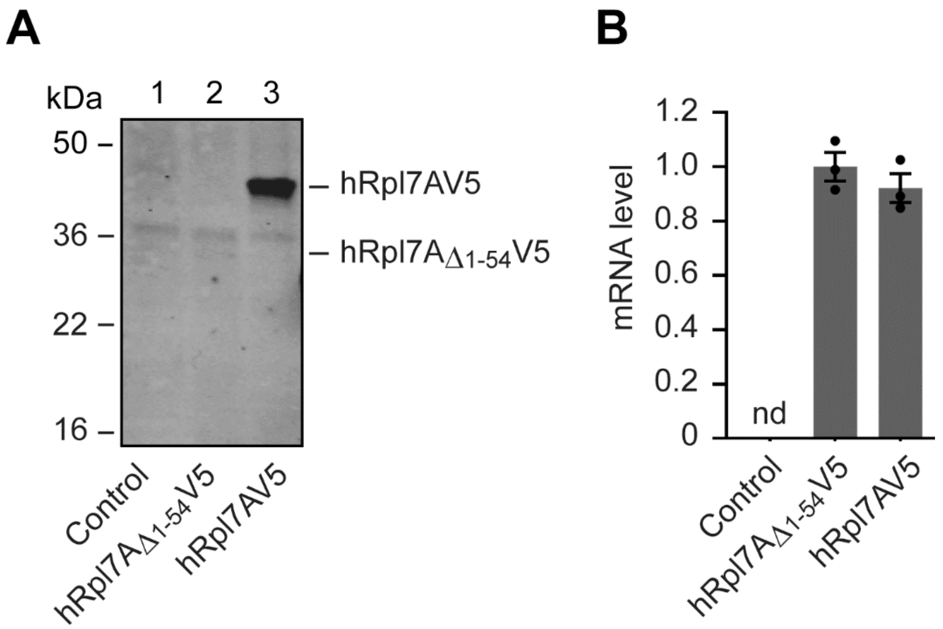


**Figure S6. The N-terminal IDR of hRpl7A suppresses CTPD.** (A) Comparison of the steady-state levels of hRpl7AV5 and hRpl7A$_{\Delta 1\text{-}54}$V5 by Western blotting with anti-V5 antibody. C-terminally V5-tagged hRpl7A and a mutant deleted of the first 54 amino acids (hRpl7A$_{\Delta 1\text{-}54}$) were expressed in HEK293 cells. (B) qRT-PCR analysis of the transcripts encoding hRpl7AV5 and hRpl7A$_{\Delta 1\text{-}54}$V5. HEK293 cells with mock transfection served as a negative control. PCR was performed with primers corresponding to the V5 tag sequence and the BGH reverse priming site on vector pEF6/V5-His A. U6 was used as an internal control for normalization of gene expression. Data are presented as mean ± S.D. from three independent experiments.

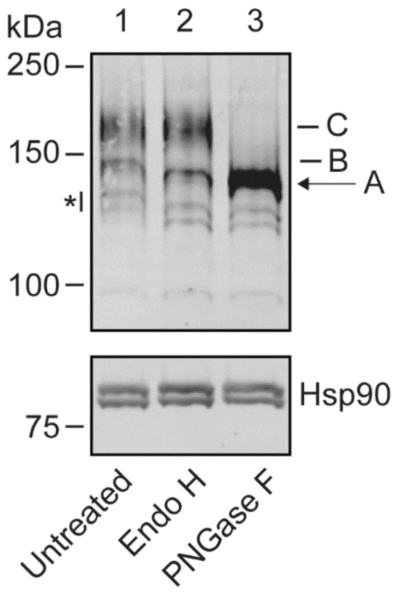


**Figure S7. Deglycosylation of CFTR proteins.** Cell lysate containing hRpl7A$_{1\text{-}55}$-CFTRV5 was untreated (lane 1) or treated with Endo H (lane 2) or PNGase F (lane 3), followed by Western blotting with anti-V5 and anti-Hsp90 antibodies. Endo H converted core-glycosylated hRpl7A$_{1\text{-}55}$-CFTRV5 (band B) but not the complex-glycosylated species (band C) to non-glycosylated form (band A). PNGase F was able to convert both band B and band C to band A. The cleavage products marked by an asterisk were also deglycosylated.

**Table S1.** Plasmids used in this study.

| Plasmid | Source |
|---|---|
| pRS314CUP1DHFRha | Ref. 30 |
| pRS314CUP1RPL8A$_{1-100}$-DHFRha | This study |
| pRS314CUP1RPL8A$_{1-50}$-DHFRha | This study |
| pRS314CUP1RPL8A$_{51-100}$-DHFRha | This study |
| pRS314CUP1RPL8A$_{1-25}$-DHFRha | This study |
| pRS314CUP1RPL8A$_{26-50}$-DHFRha | This study |
| pRS314CUP1UB-DHFRha | Ref. 30 |
| pRS313CUP1RPS31ha | Ref. 25 |
| pRS313CUP1RPL8A$_{1-50}$-RPS31ha | This study |
| pRS313CUP1UB-RPS31ha | Ref. 25 |
| pRS314CUP1RPL8Aha | Ref. 25 |
| pRS314CUP1RPL8A$_{\Delta 1-50}$ha | This study |
| pRS314CUP1RPS12ha | Ref. 25 |
| pRS314CUP1UB-RPS12ha | Ref. 25 |
| pRS314CUP1RPL8A$_{1-50}$-RPS12ha | This study |
| pRS314CUP1GFP | This study |
| pRS314CUP1RPL8A$_{1-50}$-GFP | This study |
| pRS314CUP1UB-GFP | This study |
| pRS314CUP1RPN4$_{172-229}$-DHFRha | This study |
| pRS314CUP1RPL8A$_{1-50}$-RPN4$_{172-229}$-DHFRha | This study |
| pRS314CUP1UB-RPN4$_{172-229}$-DHFRha | This study |
| pRS426CUP1GSTFlag | This study |
| pRS426CUP1RPL8A$_{1-50}$-GSTFlag | This study |
| 314CUP1APE2$_{1-10}$-DHFRha | This study |
| 314CUP1SPN1$_{1-10}$-DHFRha | This study |
| 314CUP1ERO1$_{1-10}$-DHFRha | This study |
| 314CUP1PUT2$_{1-10}$-DHFRha | This study |
| 314CUP1BGL2$_{1-10}$-DHFRha | This study |
| 314CUP1SSO2$_{1-10}$-DHFRha | This study |
| 314CUP1HRK1$_{1-10}$-DHFRha | This study |
| 314CUP1MRPS9$_{1-10}$-DHFRha | This study |
| 314CUP1RCK2$_{1-10}$-DHFRha | This study |
| pEF6hRPL7AV5 | This study |
| pEF6hRPL7A$_{\Delta 1-54}$V5 | This study |
| pEF6CFTRV5 | This study |
| pEF6hRPL7A$_{1-55}$-CFTRV5 | This study |

**Table S2. List of transcripts analyzed by qRT-PCR and the primers used**

| Transcript | Upstream primer | Downstream primer |
|---|---|---|
| *DHFRha* | YX1084: TTCCCAGAAATTGATTTGGG | YX1018: CCAGAGCTCCTAGGCGTAATCTGGGACATCG |
| *RPL8A$_{1-100}$-DHFRha* | YX1084: TTCCCAGAAATTGATTTGGG | YX1018: CCAGAGCTCCTAGGCGTAATCTGGGACATCG |
| *RPL8A$_{1-50}$-DHFRha* | YX1084: TTCCCAGAAATTGATTTGGG | YX1018: CCAGAGCTCCTAGGCGTAATCTGGGACATCG |
| *RPL8A$_{51-100}$-DHFRha* | YX1084: TTCCCAGAAATTGATTTGGG | YX1018: CCAGAGCTCCTAGGCGTAATCTGGGACATCG |
| *UB-DHFRha* | YX1084: TTCCCAGAAATTGATTTGGG | YX1018: CCAGAGCTCCTAGGCGTAATCTGGGACATCG |
| *RPS31ha* | YX1014: GAAGATCAAGCACAAGCA | YX1018: CCAGAGCTCCTAGGCGTAATCTGGGACATCG |
| *UB-RPS31ha* | YX1014: GAAGATCAAGCACAAGCA | YX1018: CCAGAGCTCCTAGGCGTAATCTGGGACATCG |
| *RPL8A$_{1-50}$-RPS31ha* | YX1014: GAAGATCAAGCACAAGCA | YX1018: CCAGAGCTCCTAGGCGTAATCTGGGACATCG |
| *RPL8Aha* | YX1034: CACGTCGACACTGAAGTCAGAGCCGAAGAC | YX1018: CCAGAGCTCCTAGGCGTAATCTGGGACATCG |
| *RPL8A$_{\Delta 1-50}$ha* | YX1034: CACGTCGACACTGAAGTCAGAGCCGAAGAC | YX1018: CCAGAGCTCCTAGGCGTAATCTGGGACATCG |
| *RPN4$_{172-229}$-DHFRha* | YX1084: TTCCCAGAAATTGATTTGGG | YX1018: CCAGAGCTCCTAGGCGTAATCTGGGACATCG |
| *RPL8A$_{1-50}$-RPN4$_{172-229}$-DHFRha* | YX1084: TTCCCAGAAATTGATTTGGG | YX1018: CCAGAGCTCCTAGGCGTAATCTGGGACATCG |
| *UB-RPN4$_{172-229}$-DHFRha* | YX1084: TTCCCAGAAATTGATTTGGG | YX1018: CCAGAGCTCCTAGGCGTAATCTGGGACATCG |
| *hRPL7AV5* | YX1104: GGTAAGCCTATCCCTAACCCT | YX1105: TAGAAGGCACAGTCGAGG |
| *hRPL7A$_{\Delta 1-50}$V5* | YX1104: GGTAAGCCTATCCCTAACCCT | YX1105: TAGAAGGCACAGTCGAGG |
| *CFTRV5* | YX1104: GGTAAGCCTATCCCTAACCCT | YX1105: TAGAAGGCACAGTCGAGG |
| *hRPL7A$_{1-55}$-CFTRV5* | YX1104: GGTAAGCCTATCCCTAACCCT | YX1105: TAGAAGGCACAGTCGAGG |
| *U6* | U6-F: CTCGCTTCGGCAGCACA | U6-R: AACGCTTCACGAATTTGCGT |
| *ACT1* | YX814: TCCATCCAAGCCGTTTTG | YX815: CGGCCAAATCGATTCTCA |
